# Machine learning analysis of naïve B-cell receptor repertoires stratifies celiac disease patients and controls

**DOI:** 10.1101/2020.11.09.371336

**Authors:** Or Shemesh, Pazit Polak, Knut E.A. Lundin, Ludvig M. Sollid, Gur Yaari

## Abstract

Celiac disease (CeD) is a common autoimmune disorder caused by an abnormal immune response to dietary gluten proteins. The disease has high heritability. HLA is the major susceptibility factor, and the HLA effect is mediated via presentation of deamidated gluten peptides by disease-associated HLA-DQ variants to CD4+ T cells. In addition to gluten-specific CD4+ T cells the patients have antibodies to transglutaminase 2 (autoantigen) and deamidated gluten peptides. These disease-specific antibodies recognize defined epitopes and they display common usage of specific heavy and light chains across patients. Interactions between T cells and B cells are likely central in the pathogenesis, but how the repertoires of naïve T and B cells relate to the pathogenic effector cells is unexplored. To this end, we applied machine learning classification models to naïve B cell receptor (BCR) repertoires from CeD patients and healthy controls. Strikingly, we obtained a promising classification performance with an F1 score of 85%. Clusters of heavy and light chain sequences were inferred and used as features for the model, and signatures associated with the disease were then characterized. These signatures included amino acid (AA) 3-mers with distinct bio-physiochemical characteristics and enriched V and J genes. We found that CeD-associated clusters can be identified and that common motifs can be characterized from naïve BCR repertoires. The results may indicate a genetic influence by BCR encoding genes in CeD. Analysis of naïve BCRs as presented here may become an important part of assessing the risk of individuals to develop CeD. Our model demonstrates the potential of using BCR repertoires and in particular, naïve BCR repertoires, as disease susceptibility markers.

## 1 INTRODUCTION

The adaptive immune system includes an extremely large repertoire of diverse lymphocyte receptors that can bind to any antigen it encounters.^1^ The high diversity of B cell receptors (BCRs) and antibody repertoires constrains our ability to measure and analyze them.^2^ Hence, what can be learned from deep immunoglobulin sequencing is highly dependent upon the sample preparation and statistical analysis utilized.^3,4^ BCRs have recently been investigated by several innovative approaches, including artificial intelligence (AI) techniques. AI is leveraged to start deciphering how information is encoded in adaptive immune receptor repertoires. In particular, there is increasing interest in applying the pattern recognition capacity of machine learning (ML) to decoding the adaptive immune receptor encoded information and using it for classification and prediction in a wide variety of applications.^5^ It includes predicting the presence of diseases, immunity status post-vaccination or infection, and designing antibody-based therapeutics.^6–10^

Upon stimulation of the adaptive immune system with antigen, the antigen-specific T- and B-cells are recruited from the repertoire of naïve T and B cells.^11,12^ Thus, the composition of the naïve T- and B-cell repertoires influences the response to the antigen. Recent work has demonstrated that an individual’s naïve T- and B-cell repertoires are shaped by genetically-determined biases.^13–15^ Thus, to analyze how the composition of naïve T- and B-cell repertoires is genetically influenced in immune disorders is of interest.

CeD is particularly interesting due to its high heritability and prevalence.^16^ Currently the only available treatment for the disorder affecting about 1% of the general population is a life-long gluten-free diet. Prevention of disease development in at risk individuals is an important goal. A recent report indicated that introduction of gluten from age 4 months is associated with reduced CeD prevalence, suggesting that disease prevention by dietary intervention may be achievable^17^

CeD is an autoimmune disorder that develops as a consequence of an inappropriate immune response to dietary gluten proteins.^18^ Typically, CeD patients consuming gluten have serum antibodies (IgA, IgG, IgM) to the autoantigen transglutaminase 2 (TG2)^19,20^ and to deamidated gluten peptides (DGP).^21–23^ Serology detecting such antibodies plays an important role in the diagnostic work-up of the disease. CeD is a polygenic disorder. Genome wide association studies have implicated 43 predisposing loci that collectively explain some 50% of the genetic variance in CeD.^24^ HLA is by far the chief genetic determinant. *HLA-DQA1* and *HLA-DQB1* genes that encode the HLA-DQ2.5, HLA-DQ2.2 and HLA-DQ8 allotypes are responsible for this HLA effect.^25^ Indeed, almost all patients with CeD carry one of these HLA-DQ allotypes. The HLA-DQ allotypes are also prevalent in the general population. Thus, HLA is a necessary but not sufficient factor for CeD development. In the genome wide association studies, no signals were reported for TCR or BCR encoding loci. Importantly, however, the coverage for these loci in the genotyping arrays used was sparse,^26^ leaving the impact of BCR and TCR encoding loci on CeD susceptibility unclear.

Here, we investigated whether naïve BCR repertoires are particular to patients with CeD. We applied an ML approach to analyze a data set of CeD and healthy subjects, and found that these clinical groups can be well classified. We further characterized motifs that could be used as biomarkers for CeD, and demonstrated that CeD patients, in contrast to healthy controls, develop clusters of BCRs with distinct properties. This study may have clinical implications for identification of individuals at risk for CeD, and provide insight into immunological processes involved in the onset of CeD and autoimmunity in general.

## 2 MATERIALS AND METHODS

### 2.1 Data

The repertoires composing the dataset were collected from blood samples of one hundred Norwegian human donors, 52 individuals with CeD and 48 healthy controls. Heavy and light BCR chains were sequenced from sorted naïve B cells. CeD was diagnosed according to standard guidelines.^27^ The research is covered by the approval of the Regional Ethical Committee (projects REK 2010/2720 and REK 2011/2472, project leader Knut E. A. Lundin). Naïve B cells were sorted on a FACSAria flow cytometer (BD) by including cells with CD19+, CD27-, IgD+, IgA- and IgG-. cDNA libraries of heavy and light chains were constructed using a 5’RACE with unique molecular identifiers (UMIs) protocol. De-identified sequence data was sent to Gur Yaari’s lab for analysis. More details about the experimental protocol can be found in.^28^

### 2.2 Preliminary Processing of Data

Raw reads were submitted to a preliminary processing pipeline as previously described.^28^ pRESTO^29^ was applied to produce high-fidelity repertoires. Sequences were then aligned to V, D, and J alleles using IgBLAST.^30^ This was followed by additional exclusion criteria: (a) Sequences whose CDR3 length is not a multiple of three (i.e. cannot be translated into AAs). (b) Sequences with isotypes other than IgM/IgD. (c) Samples with low sequencing depth after filtration (< 2000 reads). For each individual, new alleles followed by a personalized genotype were inferred using TIgGER,^31,32^ following the pipeline described in.^33^ After all filtrations, the datasets included 92 samples: 48 individuals with CeD and 44 healthy controls.

### 2.3 Repertoire Representation

Three types of repertoire representations were constructed, and were then used as features to train an ML model. This included: (1) sequence annotation-based features, (2) physicochemical properties of AA snippets, and (3) sequence similarity clusters.

#### 2.3.1 Sequence Annotation-based Features

There are common useful sequence annotation types that can be attributed using a sequence alignment process.^34^ These annotations were used to represent repertoires by a single matrix that includes the annotation frequency measurements, such as gene usage, the occurrence of combinatorial joining of genes as well as properties derived from the sequence composition. Supplementary Table S1 lists examples of the extracted sequence annotation features.

#### 2.3.2 Physicochemical Properties

It is reasonable to assume that in specific regions that are important for binding affinity, molecular characteristics are conserved. To identify these regions, we formed a representation of BCR sequences that describes the basic properties of AAs of the third complementarity determining region (CDR3). We focused on CDR3 because it provides the most important variable sequence for antibody specificity.^35^ To regulate the variable lengths of CDR3s, each CDR3 sequence was broken down into overlapping subsequences of equal length (i.e., AA k-mers). The properties of these AAs were calculated using five Atchley factors, as described in,^7^ using Change-O.^36^ Each of the five factors roughly corresponds to several general biological characteristics: polarity, propensity for secondary structure, molecular size, codon diversity, and electrostatic charge.

#### 2.3.3 Sequence Similarity Clusters

Functional V(D)J sequences from each subject were assigned into clusters based on the following criteria: (a) Having the same CDR3 length, V and J gene segment assignments. (b) The Hamming distance between the AAs of each pair of CDR3s does not exceed 0.15 (i.e. maximal dissimilarity between any two CDR3 sequences in a cluster never exceeds 15%). We applied this clustering approach using three distance thresholds:T=0.15, T=0.25 and T=0.6 under the complete-linkage condition. To avoid irrelevance or biases, clusters with fewer than N independent members (per threshold: T=0.15, N=7; T=0.25, N=10;T=0.6, N=20) were removed. The consensus sequence of a cluster is defined by a hypothetical center sequence that minimizes the sum of distances to all sequences in the cluster.

### 2.4 Machine Learning Process and Feature Selection

For a given prediction task, there may be features that do not correlate well with the response variable. To identify the features that optimize the prediction performance, we used a feature selection step. Ranking and selection were applied by the Random Forest (RF) algorithm. The RF algorithm consisted of one hundred decision trees and the result was an average of ten-fold cross-validation (over the training set that comprises 80% of the samples). Having derived a subset of the one thousand most important features, we used the Logistic Regression-Recursive Feature Elimination algorithm (LR-RFE)^37,38^ to select the strongest 300 predictor features. This number was determined by using the RFE cross-validation (RFECV) algorithm. RFECV is also a function of the sklearn python library.^39^ This function performs cross-validated selection of the optimal number of features, by removing 0 to N features using recursive feature elimination, then selecting the best subset based on the cross-validation F1-score of the model. To assess model performance, we performed 20% samples holdout cross-validation, where the model is trained on the remaining 80% of data and then is scored based on the holdout samples after selection of the best stratification features (Figure 1). This provides an unbiased evaluation and avoids the possibility of over-fitting. The model initially normalizes the training set using Z-score, and uses the training parameters to scale the holdout data. We applied this approach to repertoires, which were represented by sequence annotation features and clusters features. From the training subset, clusters were selected by having a minimum of 70% members from a particular group, celiac or healthy. The model was performed for each of the heavy and light chain datasets separately and for a combination of both.

**Fig 1.**
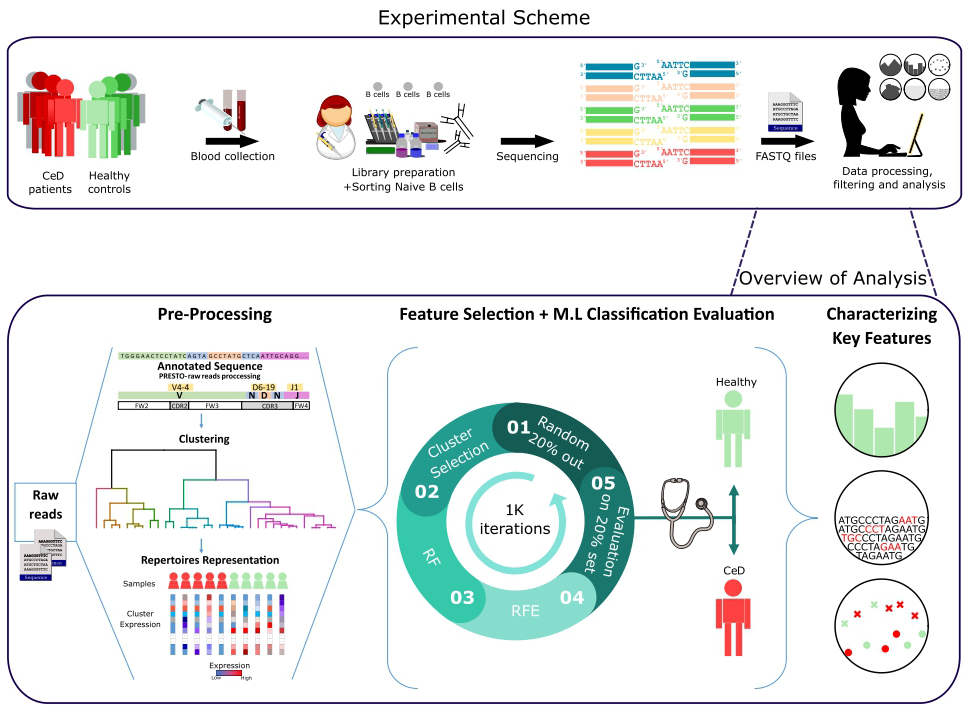
Workflow scheme. Experimental and computational analysis included the following steps: collection of blood samples from individuals with CeD and healthy controls, sequencing of naïve B cell repertoires, and analyzing the repertoires. Repertoire analysis consisted of V(D)J sequence annotation, creation of repertoire representations by antibody clustering, identifying clinical-predictable features using ML methods, and characterizing disease biomarker motifs.

### 2.5 Sequence-Based Classification

To classify antibody repertoires to their associated clinical condition based on the CDR3 AA sequences, a multiple instance learning (MIL) method with LR prediction function was used as described in.^7^ This method uses the biophysiochemaical properties (Atchley factors) of AA k-mers as the model’s features.

### 2.6 Feature Ranking

For further analysis, we needed to infer the relative importance of the different features for classification. Ranking features was done by subsampling in combination with selection algorithms. Here we ranked the features based on the recursive feature elimination (RFE) results with 1K subsampling sets. The idea was to apply a feature selection algorithm to different subsets of data and with different subsets of features. After repeating the process a number of times, the selection results can be aggregated, for example by checking how many times a feature ended up being selected as important when it was in an inspected feature subset. If the results are consistent across the subsets, it is relatively safe to trust the stability of the method on this particular data and therefore straightforward to interpret the data in terms of ranking (the more a feature is selected, the greater its importance).

### 2.7 Association between Features and Clinical Conditions

To reveal which features (predictors) are associated with the decision of a positive (CeD) or negative (healthy controls) classification, we looked at the summary of LR weights over all subsamples. As described, LR is chosen as the final classification scheme of our model. Intuitive understanding of the “influence” of a given parameter in an LR classification model, may be obtained from the magnitude and the direction of its coefficient. The weighted sum in logistic function indicates the classification probability by the following formula:

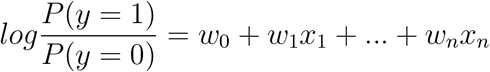

Where the term in the log() function is the probability of an event (CeD in this case) divided by probability of no event (controls), and *w_i_* is the weight for the feature value *x_i_*. Thus, in our case, a positive coefficient value means that this feature increased the probability of a CeD categorization, and a negative value indicates that this feature increased the probability of the control categorization. A highly ranked feature with a positive weight summary value was considered as a CeD-associated feature.

### 2.8 Cluster Set Enrichment Analysis (CSEA)

CSEA is used to identify motifs enriched in ranked cluster lists, where ordering is based on a measure of clusters-features importance. CSEA identifies whether members of a cluster set S tend to occur toward the top or bottom of a ranked list L, in which case the cluster set is defined by the V-gene/J-gene/CDR3-length class (i.e, clusters sharing the same V-gene category are defined to be in set *S*). This analysis is a derivative of a published analysis called GSEA.^40^ CSEA steps:

1. Divide the N clusters to sets according to attribute D (in our case V-gene/J-gene/CDR3-length group).
2. Order the N clusters by their importance values, from the maximum to the minimum (the ordered list is denoted by L). Where the rank of the i-th cluster is denoted by *t_i_*.
3. Compute the Enrichment Score (ES): start with ES = 0; walk down the ranked list L, from the top rank (i=1) to the bottom rank (i=N), increasing ES by |*r_i_*|/Σ_*j∈S*_|*r_j_*| if the i-th cluster belongs to the cluster set S, and decreasing ES by 1/(*N* – |*S*|) otherwise, where |*S*| is the number of clusters in the set S. That is, CSEA tests a null hypothesis that rankings of the clusters in a cluster set are randomly distributed over the rankings of all clusters, according to a cluster importance measure using Kolmogorov-Smirnov-like statistic.^41^
4. Take the absolute value of the maximum deviation from zero of the ES values among the N clusters as the test statistic for the cluster set S.
5. Permute the labels of attribute D and repeat steps (1)-(4). Repeat until all (or a large number of) permutations are considered.
6. Statistical significance for the association of S and D is obtained by comparing the observed value of the test statistic from (2) and its permutation distribution from (4).

The P value is computed for each cluster set by determining the empirical null distribution of enrichment scores with a permutation test procedure involving 1K data permutations. A gene set’s significance is adjusted for multiple testing by first normalizing the ES to account for the size of each cluster set, yielding a normalized enrichment score (NES). The proportion of false positives is controlled by calculating the false discovery rate (FDR) corresponding to each NES. The FDR is the estimated probability that a set with a given NES represents a false positive finding; it is computed by comparing the tails of the observed and null distributions for the NES. A cluster set was defined as significantly enriched if the false discovery rate (FDR) value was less than 0.05.

### 2.9 K-mer Enrichment Analysis

Significantly enriched k-mer motifs within the selected features were examined by counting occurrences of k-mer in sequence set that associated with the CeD group and in another sequence set corresponding control group. After the sets were defined, the sequences of both sets were broken down into overlapping subsequences (e.g., AA k-mers of length 3), and for each k-mer its frequency in the two sets was calculated. The ratio between the prevalences was used to calculate the fold change between the two sets. Then, for each k-mer we tested whether that k-mer appears significantly more times in the case or control sets, using the exact Fisher test. We controlled the FDR for multiple testing by the Benjamini–Hochberg procedure and demanded that this value is < 0.05.^42^

### 2.10 Conjoint Triad Descriptors

Conjoint triad (CT) descriptors were proposed in.^43^ In this approach, each AA sequence is represented by a vector space consisting of descriptors of AAs. To describe the properties of sequence and reduce dimensions, the twenty AA were grouped into seven classes according to their dipoles and volumes of the side chains.

## 3 Results

A graphic illustration of our overall approach is presented in Figure 1. We used two types of pre-processed BCR repertoire datasets of heavy and light chains, from individuals with CeD and healthy controls. First, feature engineering methods were used to gain different representations of the datasets and to select features that were further used to classify each repertoire to its associated clinical condition. This was followed by statistical methods for the interpretation of the model parameters.

To identify features in B-cell repertoires that are uniquely associated with CeD, we investigated which features discriminate the CeD repertoires from the healthy repertoires, using a classification model. Since the number of antibody sequences in an immune repertoire is extremely large, comparison between repertoires requires defining a smaller number of common representative features to describe each repertoire. For this, we used domain knowledge of the raw data to create features that are fit and valuable for ML modeling. To make the classification model easier to interpret and capture complex relationships, we defined sets of common representative features to describe a repertoire.

For the first repertoire representation set, we calculated the frequency of specific repertoire annotations, such as gene, allele, V-J gene combinations, and isotype usage. For the second representation set, we extracted snippets from each CDR3 and represented them by calculating the biophysicochemical properties (Atchley descriptors) of their AA sequences. Since we and others have previously reported successful immunological status classifications using IgH clusters,^44,45^ for the third set, we grouped the antibody sequences by V-J-CDR3 similarity (see methods). This essentially means that the value of each feature in the repertoire representation is the frequency of subject repertoire sequences attributed to the specific feature.

### 3.1 LR-clustering approach predicts the CeD patients based on the naïve antibody repertoire

Having derived a set of repertoire representations, we next used ML models to evaluate whether these features hold relevant information to classify individuals to CeD or to healthy based on their antibody repertoires. It is likely that most of the repertoire features are not associated with autoimmune diseases, and thus a process of feature selection is critical for the classification task. To select relevant features, we based our approach on the Random Forest (RF) algorithm and other feature-reduction algorithms, such as recursive feature elimination (RFE) and cluster selection (see methods). To determine the optimal number of features, we performed an RFE with a tuning of the number of features to select using cross-validation (RFE-CV, see methods). Logistic regression (LR) was then applied to generate the prediction on the feature-reduced dataset. We left out 20% of the samples as a test set, and trained on the remaining samples. To ensure model robustness, the process of sampling and training was repeated 1K times. An illustration of the model is shown in Figure 1.

Compared with the other repertoire representations described above, the one with CDR3 cluster features yielded a higher classification performance than those with the simple sequence annotation or Atchley factors (Supplementary Table S2). This suggests that the described repertoire annotations, including usage of specific V, D, and J genes, alleles, CDR3 lengths, biophysicochemical factors, and isotype usage, do not hold sufficient information to differentiate between the CeD and healthy groups. However, the cluster representation holds relevant information to the classification task, and the LR model with ridge (L2) regularization obtained the best classification results. Therefore, our next steps only considered the repertoire representation that is based on CDR3 sequence similarity (clusters) with LR classifier.

Next, we sought to explore the performance of repertoire classification based on the light chain dataset as well. Clusters of heavy chain (HC) and light chain (LC) sequences were inferred and used as features for the model described above. This model was applied with each dataset separately (HC, and LC sets) and with their combination (HC_LC set), using repeated cross-validation. After all filtrations, the datasets included 92 samples: 48 individuals with CeD and 44 healthy subjects. Figure 2 summarizes the average and standard deviation scores gained by the classifier on each type of dataset. The best classification was achieved using the combined HC LC datasets, where the held-out samples had an F1-score of 85%. Classification based on the HC and LC sets separately produced a lower performance, with F1-scores of 76% and 69%, respectively. To estimate the probability of correctly classifying by chance, we performed a control analysis. In the control analysis, the labels of the training data were permuted but those on the holdouts were not. The classification under these conditions seemed random, with a ~50% prediction rate.

**Fig 2.**
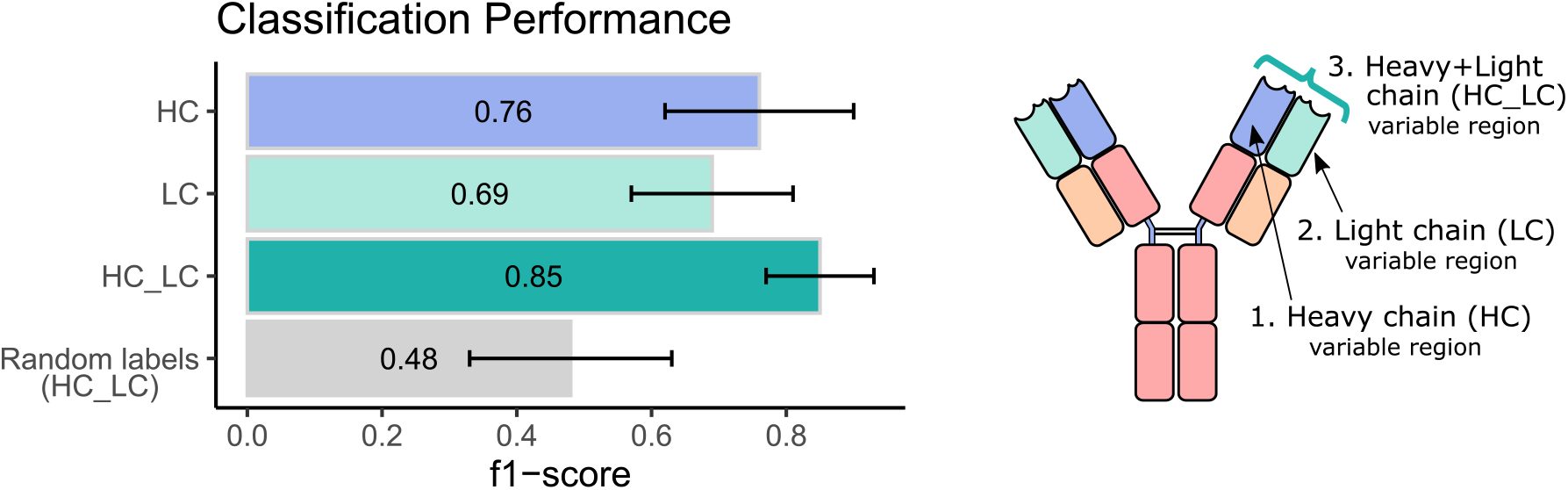
Predicting clinical diagnosis based on heavy and light chain antibody repertoire. Classification performance of the ML model using heavy and light chain variable region repertoires. Bar graphs show the average F1 score ± standard deviation (black line) across a thousand iterations. Light blue bar indicates the result using the heavy chain (HC) dataset, turquoise bar for the light chain dataset, green bar for the integrated heavy and light chain (HC_LC) dataset, and gray bar for control analysis using the HC_LC dataset with random labels.

These results indicate that there is a correlation between naïve B cell CDR3 sequence similarity clusters and CeD, and that this correlation is preserved in the repertoire representation used here. This led us to further investigate how the predictive sequences of CDR-H3, CDR-L3, and their adjacent genes vary across repertoires sampled from different clinical conditions.

### 3.2 Characterizing the key stratification features

The next step was to apply strategies for identifying components over-represented and scanning motifs, to extract patterns and knowledge from the key features that contribute to the classification performance. First, we gathered and ranked the features selected by the RFE algorithm over the 1K iterations. Of the 188 clusters that were selected in more than 40% of the iterations, 63% included genes from the IGHV3 and IGHV1 families (Figure 3B). IGHV1-69 and J4 were the highest used genes (Figure 3B, 3C), the frequencies of these genes in the overall repertoires are approximately 7% and 52%, respectively. The length distribution of CDR3 sequences, however, did not differ from a normal distribution (Figure 3D).

**Fig 3.**
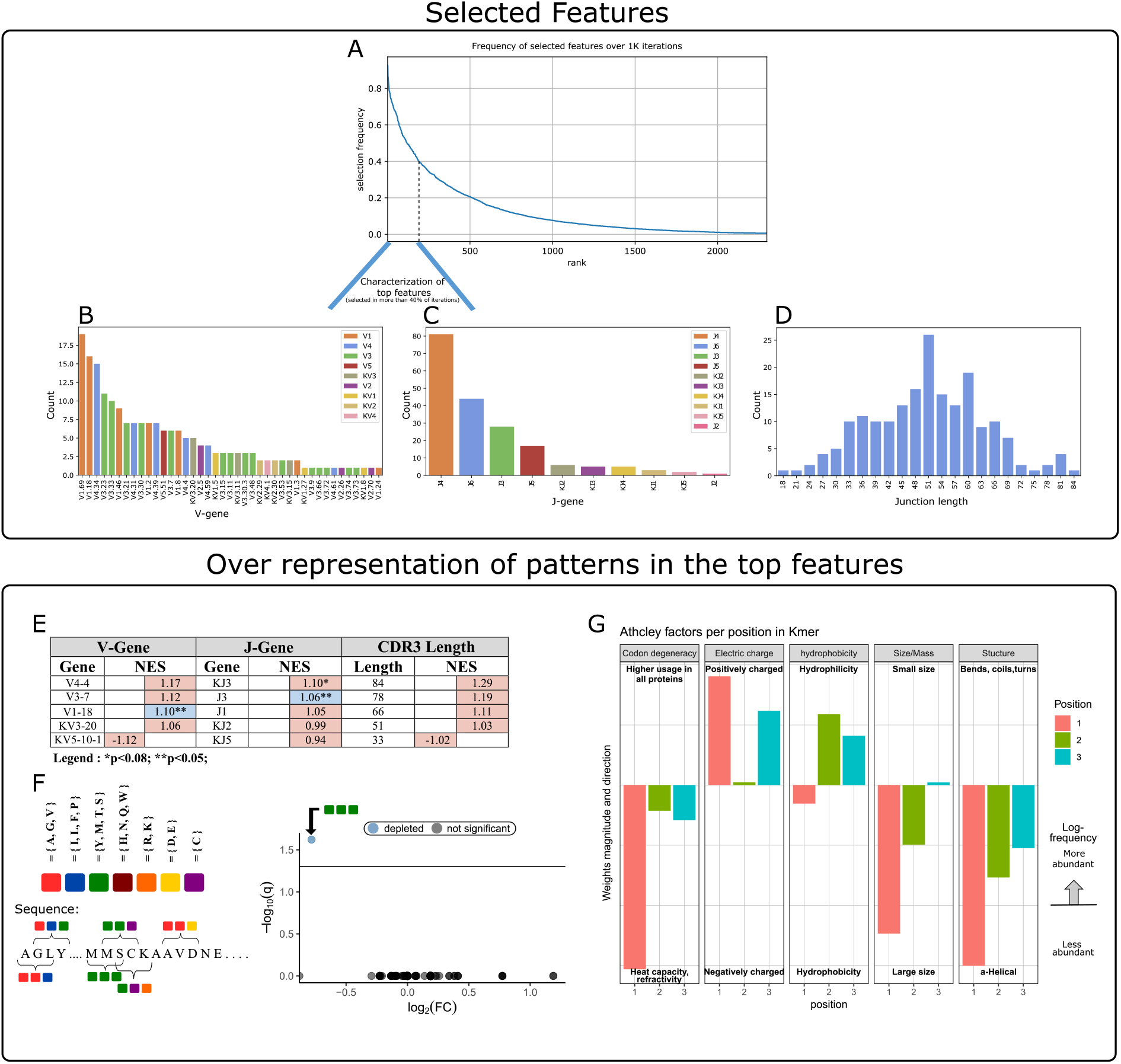
Characterization of key features used by our ML model. **(A)** Aggregation of feature selection outcomes. The Y-axis indicates the frequency of feature selection across 1K subsamples, and the X-axis represents the feature rank index. **(B)** V gene usage within the features selected in more than 40% of subsamples. **(C)** J gene usage within the features selected in more than 40% of subsamples. **(D)** CDR3 length distribution within the features selected in more than 40% of subsamples. **(E)** Enrichment cluster set results. Top normalized enrichment score (NES) outcomes of 3 cluster set analyses (V-gene, J-gene and CDR3-length). Colors indicate whether the FDR for multiple hypotheses is lower than 0.05, with blue for TRUE and red for FALSE. **(F)** Conjoint AA-triad (CT) motif is depleted in CeD-associated features. The left panel graphically displays the division of AAs to seven groups and an example representation of the possible AA-conjoint triads (as described in the methods). The right panel is a volcano plot showing the k-mer enrichment analysis with CT representation of selected clusters. The blue color indicates a point-of-interest that represents the depleted k-mer. Non-significant k-mers are shown as gray points. **(G)** The classifier weights for the three residue positions, provided by the MIL method.^7^ Columns represent the categories of five biophysicochemical factors. Positive weight values are shown as facing up bars, and negative weight values are shown as facing down bars. The length of the bar corresponds to the weight’s magnitude, and the color corresponds to the position in the snippet.

To evaluate the significance of these cluster composition results, we used cluster set enrichment analysis (CSEA) as described in the methods section. In this case, the cluster set correlated with the V-gene/J-gene/CDR3-length class, i.e., clusters sharing the same V-gene category. CSEA tests a null hypothesis in which rankings of the clusters in a set are randomly distributed over the rankings of all clusters. We observed that the IGHV1-18 and IGHJ3 cluster sets were significantly upregulated (FDR< 0.05), and the IGKJ3 set was also upregulated with FDR< 0.08 (Figure 3E).

We next sought sequence motifs that are enriched in clusters associated with CeD or healthy controls and explored their properties. For this, we used k-mer enrichment analysis (see methods) to compare the case set of sequences (100 top clusters associated with the CeD cohort) to the control set of sequences (100 top clusters associated with the healthy controls). To avoid biases caused by unbalanced sets, an equal distribution of sequence lengths from each set was sampled, without replacement. Here, cluster clinical association was determined by the LR coefficients, and clusters were represented by their CDR3 consensus sequence, as described in the methods. To provide the optimal k-mer, we examined different lengths and types of k-mer representations (K=2/3/4/5, type=nucleic/AA/AA properties). The k-mer representation that showed significant results was provided by conjoint AA-triad (CT) properties descriptors, in which residue-specific physicochemical interaction effects were taken into account. Figure 3F shows a highly significant result for one of the CT types, by a blue data point appearing at the top left of the volcano plot, meaning that we identified an AA-triad that is depleted in clusters associated with CeD versus healthy controls. This CT type is composed of three AAs that belong to the same class {Y’M’T,S}. The dipoles and volumes of these AAs, side chains are characterized by 1.0 < Dipole < 2.0 (*Debye*) and Volume> 50 (*Angstrom*^3^), respectively.

Another motif that we have scanned is a biophysicochemical feature in the CDR3 sequences of the selected clusters, taking into account the relative position of each CDR3 snippet. For this task, we applied an MIL method as presented by,^7^ to detect different 3-mer AA subsequences that share similar biophysicochemical properties in the CeD associated clusters. We have chosen this method because previous studies have reported a disease-specific biophysicochemical motif in the CDR3 of BCR or TCR chains of different types of diseases.^46^ Furthermore, this method searches for complex motifs, which may shed light on the properties of biological binding regions, and provides dimensionality reduction by extracting fewer features that hold most of the information included in the initial set. Here, the 3-mers were instances, the consensus sequences of the most highly ranked 300 clusters were bags, and the bag label was the cluster associated clinical status (150 CeD associated clusters and 150 control associated clusters). First, each consensus was divided into all the contiguous 3-mers it contains, and for each 3-mer the properties of AA residues were calculated according to five Atchley factors and its position. This model accurately classified the validation data set into associated clinical groups with 71.6% accuracy. Hence, we assume that a recurring motif can be discerned for clusters associated with CeD versus control. To reveal the pattern that associated with CeD categorization, we investigated the optimal model weights that push the classification towards a CeD categorization. Here, the probability for a CeD categorization increased for 3-mers with hydrophobic residues at the first position and hydrophilic residues at the last two positions. We also identified a preference for 3-mers with a large relative abundance of residues with large-sized alpha-helical segments and a positive charge. Thus, a cluster was classified as CeD-associated if it contained CDR3 sequences with 3-mers comprising AA residues with distinct bio-physicochemical motifs.

## 4 Discussion

In this study, we tested the ability of the naïve BCR repertoire to inform us about CeD status. We tested many repertoire features such as sequence annotation-based features and sequence similarity-based features (clusters), to develop an ML-based model to distinguish CeD patients from healthy controls. The “traditional” repertoire annotations, including usage of specific V, D, and J genes, alleles, CDR3 lengths, D sequence lengths, and isotype usage, did not remarkably differ between the CeD and healthy groups. Strikingly, however, we found that clusters with characterized common motifs can classify CeD from healthy individuals, with an F1-score of 85% for classification by naïve BCR repertoires of combined IgH and IgL sequences. Disease associated patterns were discovered and include several interesting findings. First, we observed several genes that are enriched in the prediction clusters (e.g., VH1.18 and JH3. See Figure 3E). Second, clusters that are CeD-associated are enriched for 3-mers comprising residues that share similar bio-physicochemical properties at key positions. These residues exhibit a propensity to participate in alpha-helical segments that are positively charged, hydrophilic, and relatively large (see Figure 3G).

Several underlying reasons could be responsible for the differences in naïve BCR repertories that allowed stratification of CeD patients from controls. The first one is that structural features of the naïve BCRs affect selection and survival of naïve B cells, which influences the risk for CeD development. The second is that the BCRs of naïve B cells somehow connect to the effector (i.e. antigen experienced) B cells that are involved in the development of CeD. A third possibility could be that there were impurities in the sequenced repertoires, meaning that not only naïve cells were sequenced. And finally, there could be confounding factors during cell processing or library generation that have created artifacts. It will be important to verify our results in future studies.

Anti-TG2 and anti-DGP antibodies are hallmarks of CeD, and both these antibodies across patients have stereotyped patterns of heavy and light chain usage. For anti-TG2 some of the most common combinations are IGHV5-51:IGKV1-5, IGHV3-48:IGLV5-45, IGHV4-34:IGKV1-39-pI and GHV1-69-p:IGKV1-17-p.^20,47,48^ The different heavy/light chain combinations recognize distinct conformational epitopes of TG2. For anti-DGP the most common combinations are IGHV3-23:IGLV4-69, IGHV3-15:IGKV4-1 and IGKV3-74:IGKV4-1.^22,23,49^ No signals related to these V genes were in our analysis among the identified classifiers that distinguished CeD from healthy subjects. In follow-up studies it will be interesting to investigate whether the patterns we detected in the CDR3s of CeD repertoires by ML are related to the antigen-specific antibodies that carry these gene combinations. Overall, anti-TG2 and anti-DGP antibodies have few somatic mutations, meaning the antibodies recognize their antigen in germline or near germline configuration.^20,22^ Thus, conceivably, having the “correct” V(D)J combinations in the naïve repertoire can be expected to play a role in development of the disease. The naïve CDR3 repertoire has a big element of stochastically introduced variation, which may not relate to genetics. Still, there are genetic elements within the J- and D-gene encoded sequences, and the recombination efficiency between the V, D, and J genes is certainly genetically influenced. To try and identify such explicit genetic patterns differentiating between CeD patients and controls, we included in one of the initial ML models genotype and haplotype inferences from the BCR repertoire data that were extracted by TIgGER^32^ and RAbHIT.^50^ However, these features did not increase the success of the model. Nevertheless, more complex patterns might be hidden in the regulatory regions of the BCR encoding loci.^51^ In the future, these regions may be sequenced directly using tailored long-read sequencing approaches.^52^

We explored a niche of BCR repertoire analysis that has received little to no attention in the scientific community. Although attempts to consolidate and make sense of the high-dimensional immunogenomic features that predict conditions of BCR repertoire is not a new concept, the vast majority of work of the currently available ML methods for immune receptor sequencing data have focused on the individual immune receptors in a repertoire, with the aim of, for example, predicting the antigen or antigen portion (epitope) to which these sequences bind, or to classify tumor tissue from normal tissue based on a single input sequence.^53–56^ In this work, the hypothesis is that there are shared BCR sequence patterns across individuals with a shared immune state. This principal had been used previously by us and others, for example by,^44^ who discriminated the BCR repertoires of individuals with current or past Hepatitis C virus (HCV) infection with 90% accuracy, by grouping it into clusters of BCRs.^57^ distinguished between the T cell repertoires of mice immunized with or without ovalbumin with 80% accuracy, by decomposing the TCR CDR3 sequences into overlapping AA k-mers. Several groups used ML with a feature-engineering approach based on biophysicochemical parameters of sequences, such as representing AA k-mers according to their Atchley biophysicochemical properties.^7,58–60^ These results have motivated us to apply an LR-clustering approach to classify antibody repertoires and examine motifs based on k-mers and physicochemical properties.

This work is a first step towards understanding the effect of naïve B cells on CeD development. The observations in this research can be further examined to be based on CeD-associated BCRs, like autoantibodies to TG2 and antibodies to DGP. It would also be interesting to explore the effect of IgH and IgL combination on naïve B cell populations in CeD patients. This could be examined by single-cell sequencing datasets.

Further studies of the unique autoimmune condition may provide a better understanding of the pathophysiological mechanisms involved in the pathogenesis of CeD, and possibly of other autoimmune diseases, paving the way to innovative treatment strategies. If measures to prevent CeD become a reality, the ability to identify individuals who are at risk for developing CeD will be particularly important. Analysis of naïve BCRs as presented here may become a part of this risk assessment.

## Author Contributions

G.Y. and L.M.S. conceived and supervised the project; K.L recruited study subjects; G.Y. and O.S. developed the ML approach and analyzed the data; G.Y., O.S., P.P. and L.M.S. wrote the paper. All authors edited the manuscript.

## Acknowledgments

We thank Ayelet Peres for discussions and helpful advice; We would also like to express our gratitude to all study participants.

## Data Availability Statement

The datasets analyzed for this study can be found in the European Nucleotide Archive (ENA), under accession number PRJEB26509 (ERP108501).

## Supplementary Materials

**Fig 4.**
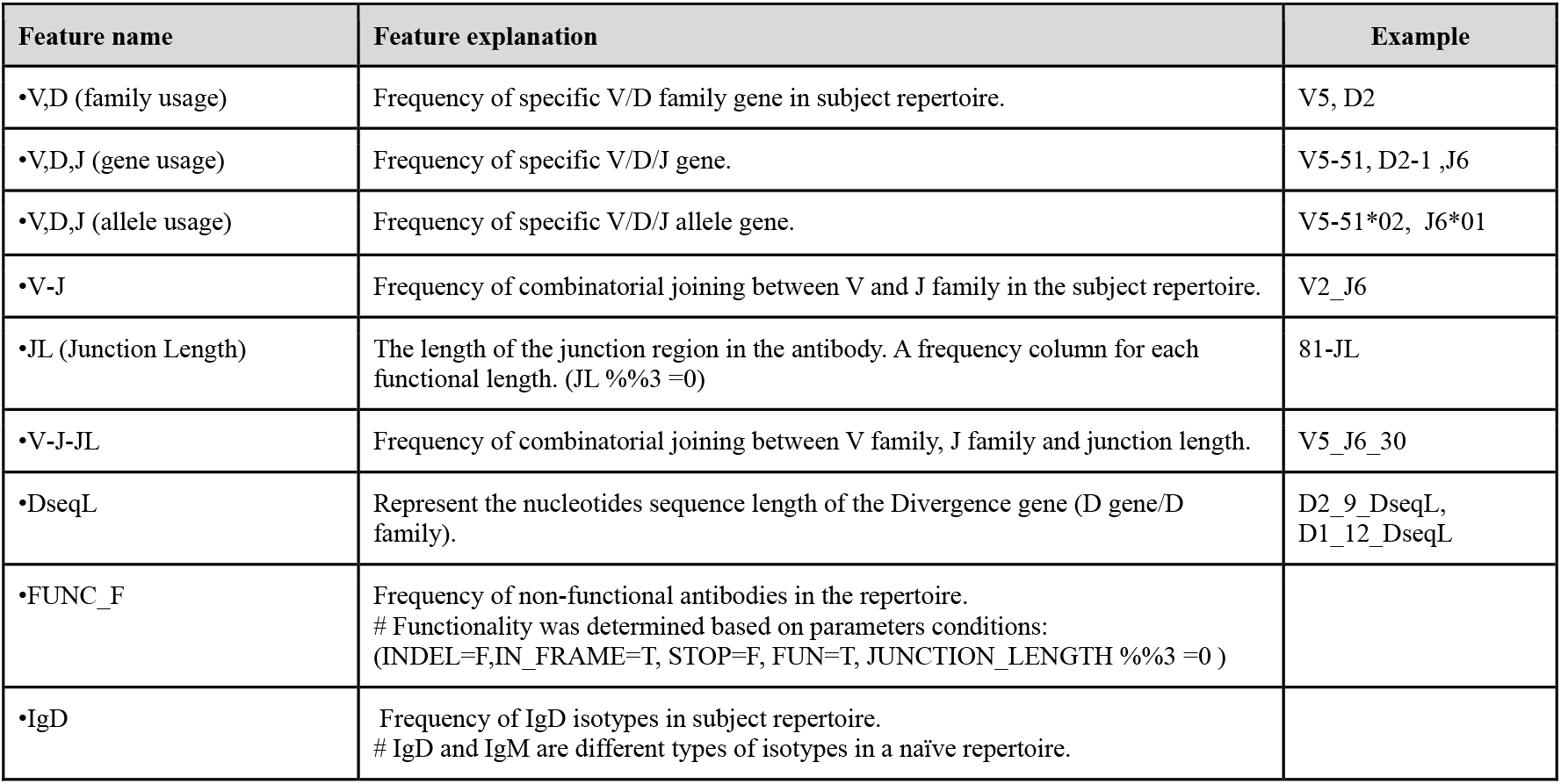
4 Sequence annotation based. This table lists examples of the extracted sequence annotation based features

**Fig 5.**
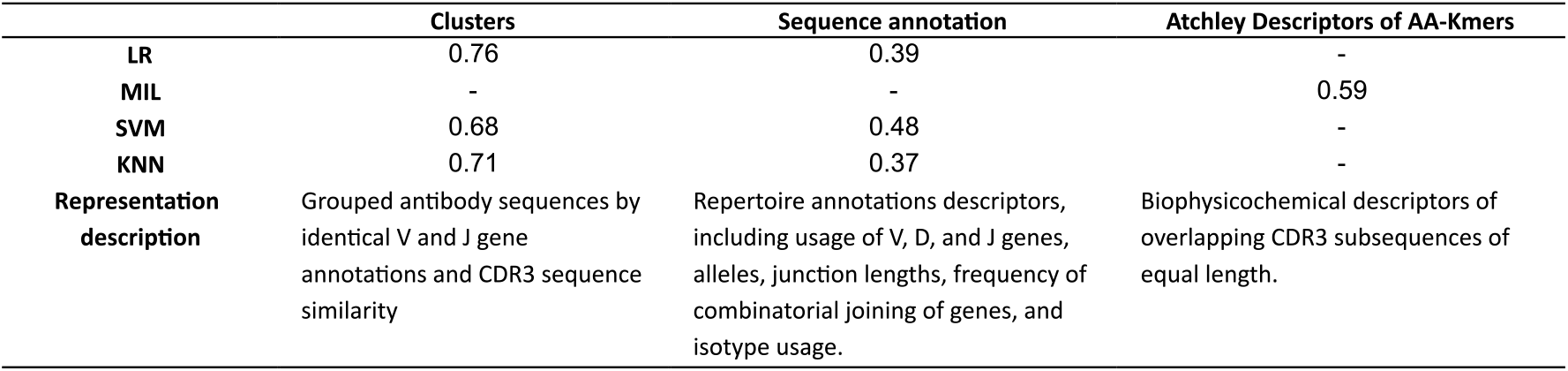
Average F1-score comparison between different models. Average F1-score comparison between different models depending on the representation of naïve antibody repertoires

